# Molecular epidemiology and clone transmission of carbapenem-resistant *Acinetobacter baumannii* in ICU rooms

**DOI:** 10.1101/2020.06.04.135624

**Authors:** Xiufeng Zhang, Fangping Li, Zhuangwei Hou, Furqan Awan, Hongye Jiang, Xiaohua Li, Zhenling Zeng, Weibiao Lv

**Author notes:** The first two authors contributed equally to this article. Correspondence: Weibiao Lv and Zhenling Zeng, and.

## Abstract

Carbapenem-resistant *Acinetobacter baumannii* (CRAB) is a major cause of nosocomial infections and hospital outbreaks worldwide, remaining a critical clinical concern. Here we characterized and investigated the phylogenetic relationships of 105 CRAB isolates on intensive care unit surfaces from one hospital in China collected over six years. All strains carried *bla_OXA-23_*, *bla_OXA-66_* genes for carbapenem resistance, also had high resistance gene, virulence factor and insertion sequences burdens. Whole-genome sequencing revealed all strains belonged to ST2, the global clone CC2. The phylogenetic analysis based on the core genome showed all isolates was dominated by a single lineage of three clusters and eight different clones. Two clones were popular during the collection time. Using chi-square test to identify the epidemiologically meaningful groupings, we found the significant difference in community structure only present in strains from separation time. The haplotype and median-joining network analysis revealed genetic differences among clusters and changes occurred overtime in the dominating cluster. Our results highlighted substantial multidrug-resistant CRAB burden in hospital ICU environment, demonstrated potential clone outbreak in hospital.

## Introduction

The spread of antibiotic-resistant bacteria poses a substantial public health crisis worldwide. *Acinetobacter baumannii*, known as a member of the ESKAPE group, is responsible for a vast array of nosocomial infections throughout the world(51). The extraordinary ability to survive in a harsh environment and to readily acquire antimicrobial resistance determinants have made *A. baumannii* as a public health threat(18). Carbapenems are known as the frontline treatment for infections multidrug-resistant *A. baumannii* infections. However, the wide spread carbapenem resistant *A. baumannii* (CRAB) strains have caused a major concern worldwide due to the limited treatment choices(13). In 2017, the World Health Organization has listed CRAB as one of the most critical pathogens and the highest priority in new antibiotic development(56).

Carbapenem resistance in *A. baumannii* is medicated by several coexisting mechanisms but the most prevalent mechanism is associated with carbapenem-hydrolyzing enzymes(9). There exist three different types of β-lactamases leading to carbapenem resistance, such as ambler class A β-lactamases (*bla*_GES-14_, *bla_TEM_, bla*_SHV_, *bla*_CTX-M_ and *bla*_KPC_), metallo-ß-lactamases (*bla*_IMP-like_, *bla*_VIM-like_, *bla*_SIM-1_, and *bla_NDM-1_*) and oxacillinases (*bla*_OXA-23-like_, *bla*_OXA-24-like_, *bla*_OXA-58-like_, *bla*_OXA-143_, *bla*_OXA-235-like_, *bla*_OXA-51-like_)(15, 52). Major expression of OXA genes might be facilitated by insertion sequence elements (ISs), such as IS*Aba1*, IS*Aba4* and IS*Aba125*, which provide an additionally strong promoter(6, 19). In addition to the resistance, CRAB also carried a wide arsenal of virulence factors predisposing for a worsening course of disease(2, 37). Although virulence determinants in *A. baumannii* are incompletely understood, the genes related to biofilm formation, outer membrane proteins, surface glycoconjugates, micronutrient acquisition systems and secretion systems were thought to important for this pathogen to successfully infect its hosts(21). Hence, improved surveillance and understanding of CRAB is a key factor in reducing death tolls caused by *A. baumannii* infections.

Numerous nosocomial outbreaks caused by CRAB have been described in the world(49, 57). In many cases, one or two epidemic strains were perceived in a certain epidemiological setting, particularly in intensive care units (ICUs)(53, 62). Molecular characterization reveals that clonal dissemination plays an important role in nosocomial outbreaks of CRAB and most of them are belonging to the global clones 1 and 2(14). This might be due to the transfer of colonized patients who transmit these epidemic strains among hospitals. To date, the international clone ST2 was the most dominant type globally of all genomes sequenced(34). Several studies showed the dissemination of CRAB isolates mainly harboring the *bla_OXA-23-like_* and belonging to ST2 in different countries(27). The recent expansion of *A. baumannii* sequenced genomes has permitted the development of large-array phylogenomic and phenotypic analyses, which can offer valuable insights into the evolution and adaptation of *A. baumannii* as a human pathogen(36, 59). Though there are many studies on the molecular mechanisms of resistance and global epidemiology of CRAB, no study was designed to investigate the distribution, molecular epidemiology, phylogenetic relationships of CRAB recovered from ICUs. Therefore, we conducted a six-year-long longitudinal study at a tertiary care hospital in China, to study the epidemiology and characterization of CRAB on healthcare surfaces using whole-genome sequencing and bioinformatic analyses.

## Materials and methods

### Strains collection and Antimicrobial susceptibility

*A. baumannii* strains from ICU rooms were collected between 2013 and 2018. Reference *A. baumannii* ATCC 19606 (strain M19606) was purchased from Guangdong culture collection center. For all isolates, the minimum inhibitory concentrations (MICs) of antibiotics were determined using a BD PhoenixTM 100 Automated Identification and Susceptibility Testing system (BD, USA) according to CLSI guideline(17).

### Illumina Whole Genome Sequencing

Bacterial genomic DNA of *A. baumannii* strains was extracted using a HiPure Bacterial DNA Kit (MAGEN). The libraries were created using the VAHTS^™^ Universal DNA Library Prep kit for Illumina. Whole genome sequencing was carried out on an Illumina Hiseq 2500 system to obtain 2×150bp reads. The sequence quality of the reads was evaluated using FastQC. Processed reads were de novo assembled into contigs with CLC Genomics Workbench 10.1 (CLC Bio, Aarhus, Denmark) and genomes were annotated by NCBI Prokaryotic Annotation Pipeline (PGAP).

### Taxonomic assignment

FastANI v1.1 (https://github.com/ParBLiSS/FastANI) with the core algorithm of BLAST-based ANI solver was used to identify the species of all isolates. Reference *A. baumannii ATCC* 19606 participated in species identification as standard sample. Species were determined if the genome in question had >95% ANI index with the type genome(48). Finally, all the isolates sequenced in this study were used to construct a Triangular matrix, representing the product of the ANI and percent genome aligned (sp1).

### ARGs, VGs, IS identification

Antibiotic resistance genes (ARGs) were annotated using the ResFinder BLAST identification program (https://cge.cbs.dtu.dk/services/ResFinder/)(30). Associated metadata was displayed as a color strip to represent bacterial isolate demographics and expected resistance to antibiotics. Virulence genes (VGs) were identified by the core database of VFDB (setA) (http://www.mgc.ac.cn/VFs/) using BLASTv2.7. ISs were annotated using IsFinder BLAST (https://www-is.biotoul.fr/index.php). The presence/absence matrix of ARGs, VGs or ISs was visualized in pheatmap R-package(R).

### Clustering and Pan-genome analysis

Prokka v1.11 was used to produce gff file for the contig of *A. baumannii* genome. After that, Roary v3.8.0 and Mafft v7.0 were used to construct a core genome alignment(28, 42). Core_genome_alignment.aln, the output file of Roary pipeline was conveyed to fastGEAR to identify instances of recombination within these samples. The recombinant regions were removed using the script of recombinant2out.py of Bacteriatool tool package (https://github.com/lipingfangs/Bacteriatool), which we designed for bacteria NGS data analysis including series of custom python scripts. Core genome alignment either removal of recombination regions and non-removal of recombination regions was used to generate a maximum likelihood tree with Iqtree v1.6.12 with 1000 times bootstraps(40). The output newick file was visualized in iTOL(33). In silico multilocus sequence typing (MLST) was performed with the MLST program. Lineages and clusters identified by hierBAPS during fastGEAR were also marked on the trees(7).

### Clonality analysis

Pairwise single nucleotide polymorphism (SNP) counts between all isolates in the recombinant corrected core genome alignment were calculated by snpdistance.py of Bacteriatool tool package. All paired distances <20 SNPs/a were considered as a clonotype to further study. Pairwise groupings were import to R to compute the euclidean distance and visualize these groupings with R package ggplot2 (http://had.co.nz/ggplot2/).

### Spatiotemporal clone linkage and population dynamics analysis

Based on SNP distance, R package vegan was used to PERMANOVA test in order to calculate the genetic diversity of the core genome between the clone types and the bayesian clusters in the sample community(44). The basic R function chisq.test() was adopted for examining the structural diversity of clonotype communities at different times and from different sources. The core genome alignment culling standard strain M19606 was conveyed to custom python script in order to extract the SNP sites which ignored site of miss. DnaSP is used for haplotype identification(50). A median joining network of haplotypes was generated by the NETWORK program to indicate the relationship between each haplotype, clone type with time development(4). DnaSP and software Arlequin were used to dynamical analyse CRAB population by establishing mismatch curve(12).

### Nucleotide Accession Number

These assemblies sequence data of *Acinetobacter baumannii* isolates were deposited in the GenBank database under BioProject accession PRJNA541408, 541366 (M19606 strain), 541386, 541822, 542047, 542048, 564419 (L9 and L40).

## Results

### ICU surfaces had high carbapenem-resistant *A. baumannii* burden

A total of 302 *A. baumannii* strains were isolated from hospital ICU rooms and 105 isolates showed resistant to imipenem and meropenem. Then we choose all these CRAB strains and reference strain *A. baumannii* ATCC 19606 for phenotypic and genomic analysis. We constructed a Hadamard matrix, which represents the product of ANI and percent of the genome aligned between every pairwise combination of the 106 genomes sequenced. Hierarchical clustering of Hadamard values confirmed 106 isolates as *A. baumannii* (Excel S1). The information about the source, year of isolation and gender of patients of all CRAB isolates were shown in Figure 1 and Excel S2. Of these, 72.3% and 75.2% of 105 CRAB strains were from sputum and from patients with pulmonary infection, respectively. Most strains (53.3%) were isolated from older people, who are more than 65 years old.

**Fig 1.**
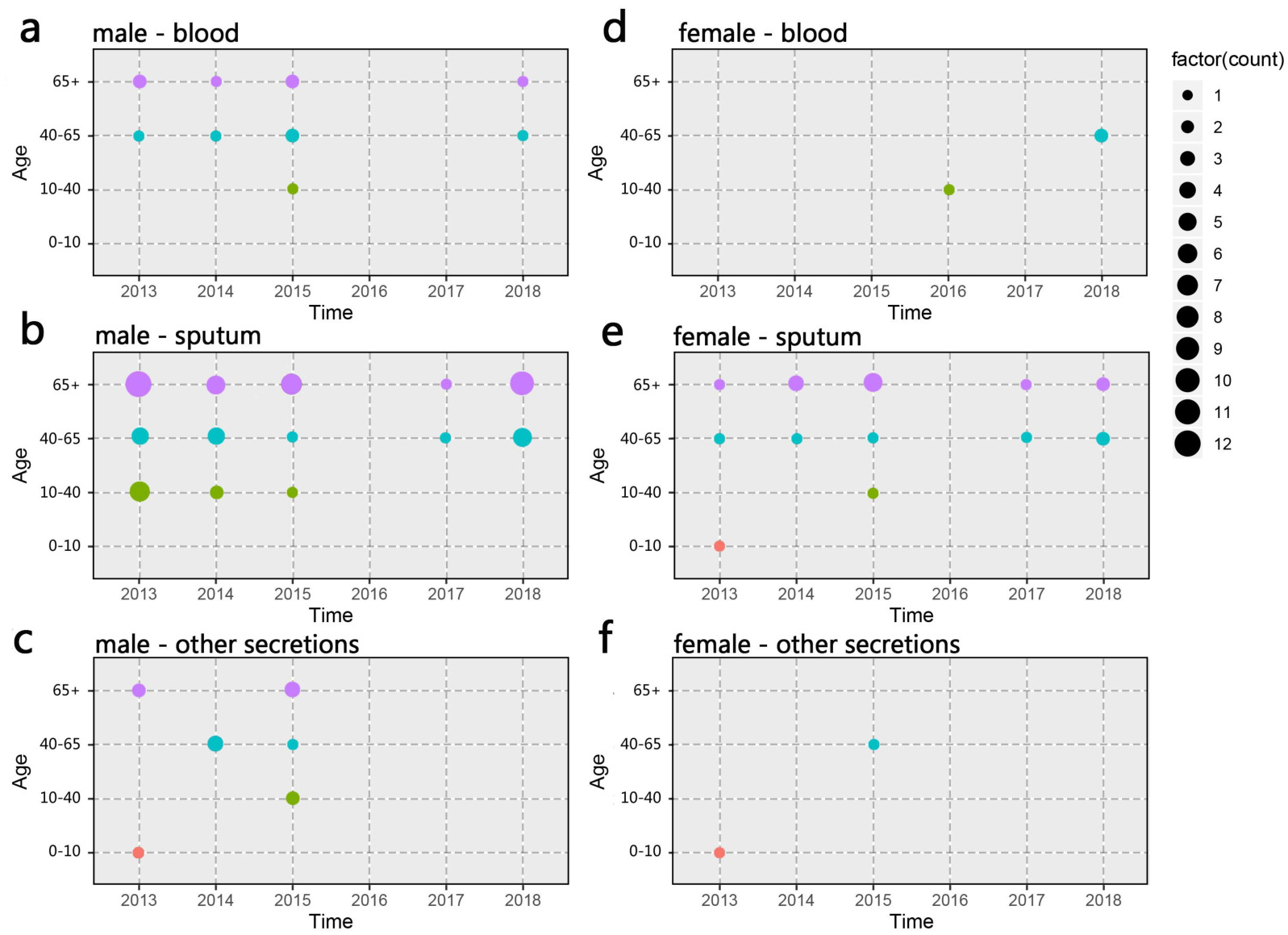
Spatiotemporal, demographic and clinical characteristics of 105 isolates of Shunde hospital. **a-c** Sample distribution from male patients; **d-f** Sample distribution from male patients; The sampling time range was 2013-2018, and the patients were divided into four groups according to age: 0-10 years old, 10-40 years old, 40-65 years old and over 65 years old. The sampling source were divided into three groups: blood, sputum and other secretions.

### Single lineages dominated CRAB population

We used Prokka to annotate protein coding sequences and constructed core genome maximum-likelihood phylogenetic trees by using Roary and RAxML. The lineages and clusters were identified by fastGEAR/BAPS.

Our results demonstrated that a single lineage represented of all CRAB strains collected over six years, which was composed of three clusters. Cluster 1, cluster 2 and cluster 3 contained 87, 17 and 1 isolates, respectively. Interestingly, all strains were assigned to ST2, which belonged to clonal complex 2 (CC2) and to international clone II. Besides, there were 9506 genes in all these strains and 99% of strains shared 3202 core genes (Fig 2). According to the core genome analysis, we found 1746 recent homologous recombination sites in core genomes of all CRAB strains and no ancestral homologous recombination sites were detected.

**Fig 2.**
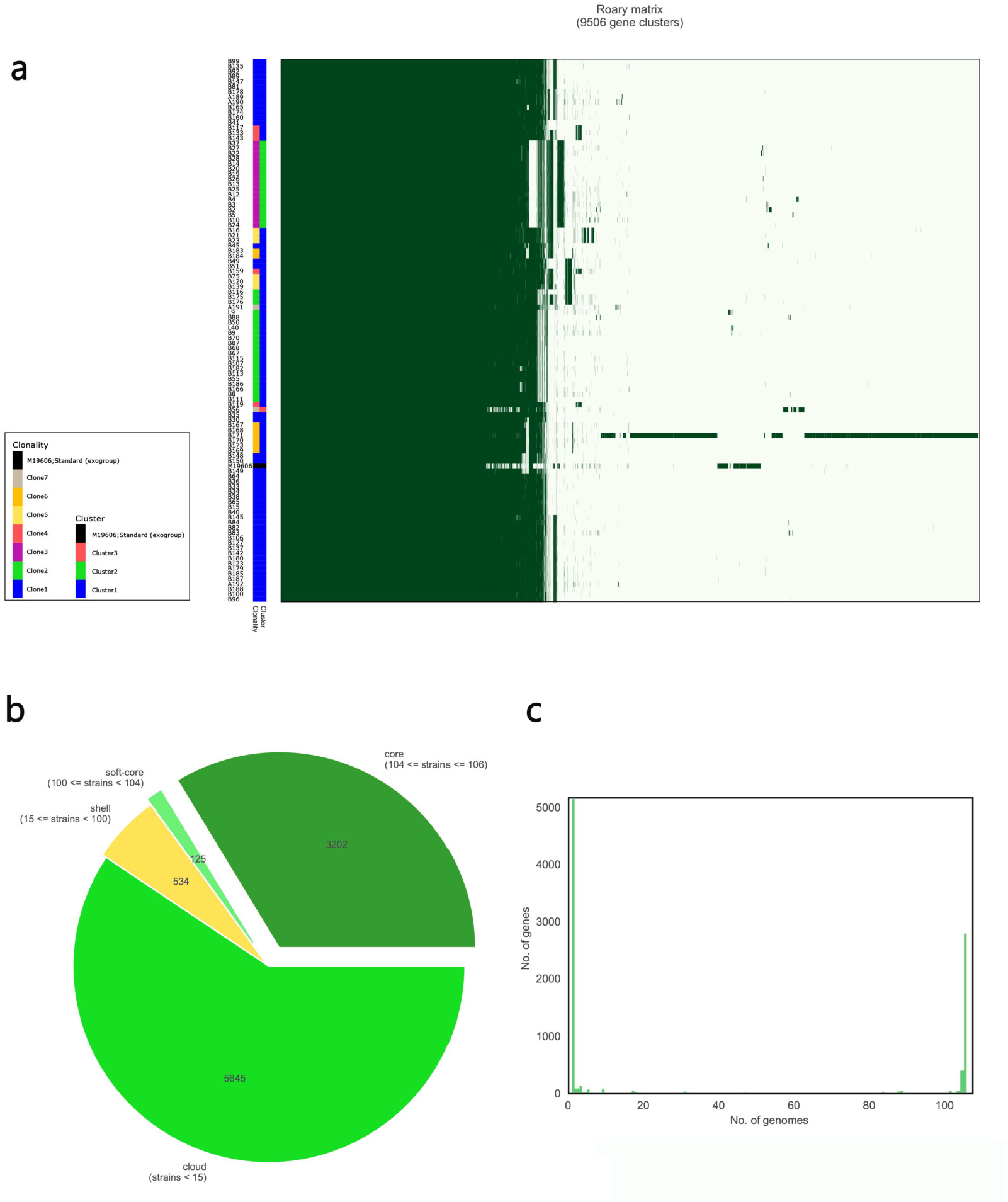
Results of pan-genomic analysis. **a** Matrix with the presence and absence of core and accessory genes; totals of 9506 genes were clustered annotation with different colors represent different clonalities based on SNP distance and different clusters based on bayesian classification. **b** Pie chart shows different kinds of accessory genes; The number of core genes was 3202. **c** Quantity distribution of accessory genes in different CRAB isoiates

### Genetically related isolates were linked by clone

To identify clone within CRAB, we removed recombinant positions from the core genome alignment and calculated pairwise SNP. Clonal groups were defined conservatively as isolates with twenty or fewer core genome SNPs in the the same species. The results showed that CRAB had multimodal pairwise SNP distance distributions, indicating concordance between SNP distances and clustering on core genome phylogenetic trees (Fig 3 and Fig 4). Eight clones (Six clones and two clone others) were observed in CRAB strains. Of these, 43.8%, 20% and 16.2% of all strains belonged to clone 1, clone 2 and clone 3, respectively. In addition, cluster 1 contained most clone groups, including clone 1, clone 2, clone 4, clone 5, clone 6 and clone other (A191), with strains collected from 2013 to 2018. Cluster 2 was only composed of clone 3, which with all the samples isolated from 2013. Cluster 3 only contained clone other (B56). The clone classification also conformed to genetic distance distribution matrix based on R package ape (Fig S1). Permanova analysis of clone groups (including clone 1, 2, 3, 4, 5, 6) based on SNP distance showed that there were extremely significant differences among different clone groups (P<0.01), which indicated that the classification is correct. However, permanova analysis between clone 1, 2, 3, 4, 5, 6 and clone others (A191 and B56) indicated that the significance difference among clones fluctuated, which is most likely due to sample size issues in clone other groups (Table 1). Similarly, permanova analysis of cluster 1 and 2 demonstrated that there were extremely significant differences among clusters (P<0.01). When cluster 3 compared to cluster 1 or cluster 2, the results of permanova analysis were also different (p=0.011 and p=0.062, respectively) (Table 2).

**Fig 3.**
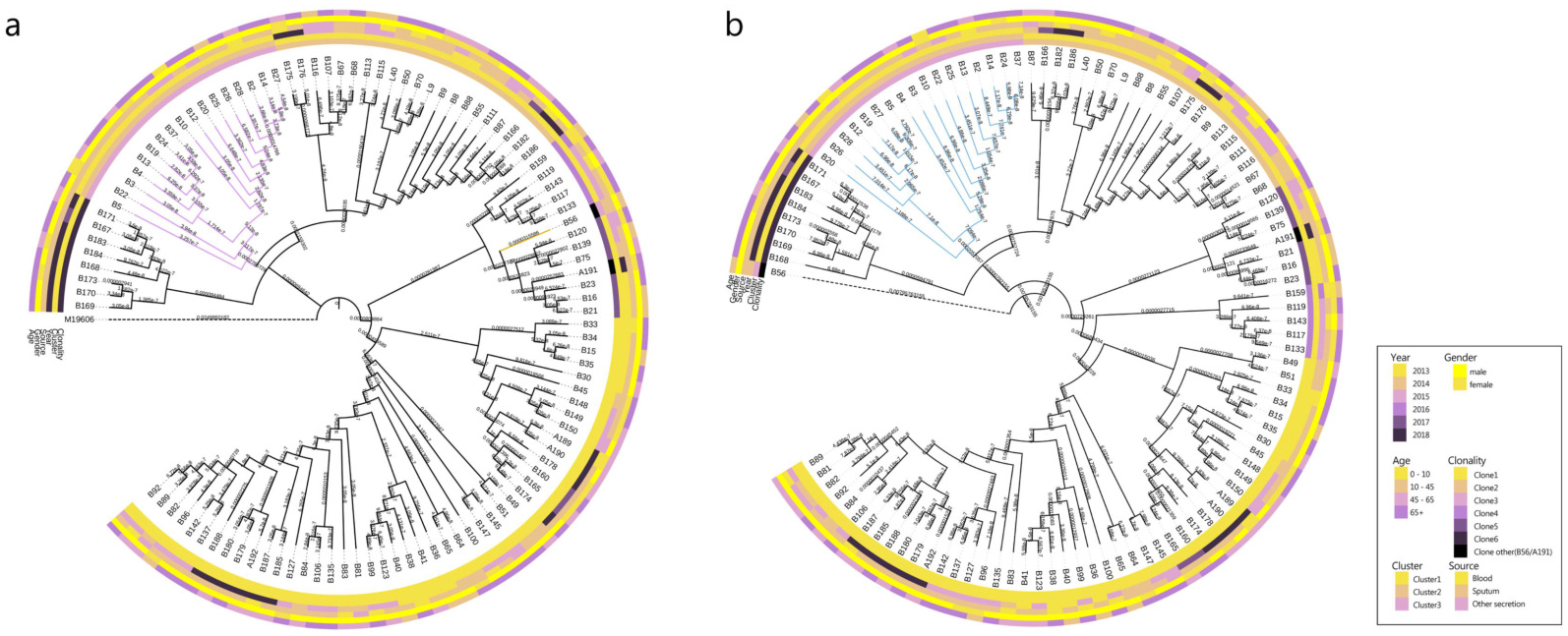
Maximum likelihood phylogenetic trees from core genome alignments of CRAB isolates. **a** Phylogenetic tree constructed from the core genomic file including the standard strain M19606 as outgroup. **b** Phylogenetic tree constructed from the core genomic file excluding he standard strain M19606; Colored annotations are added next to the main tree for sampling time and sources, patient ages and genders and cluster and clonality classification of 105 CRAB isolates.

**Fig 4.**
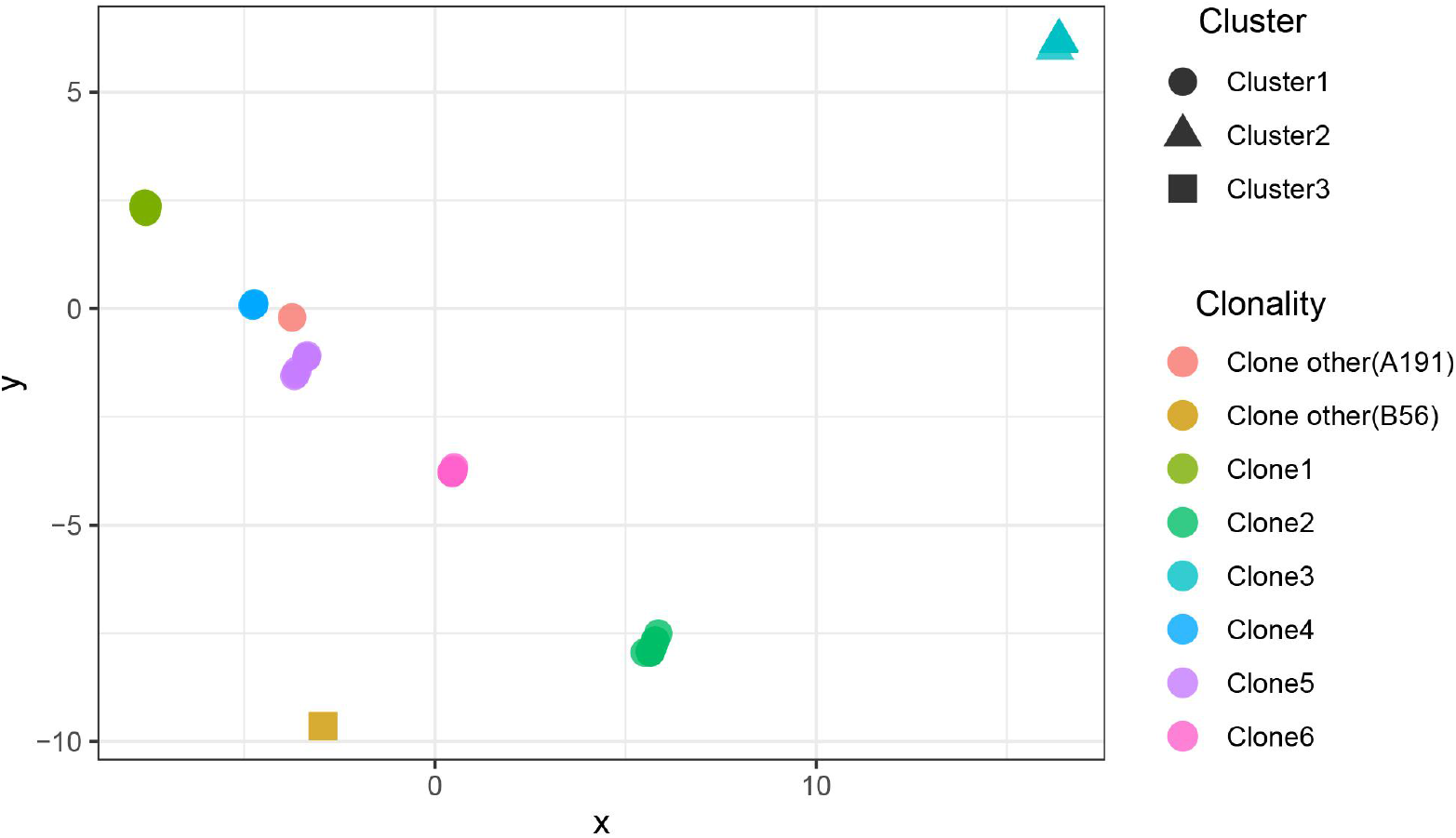
The scatter plots of SNP distance reduced dimension among CRAB isolates. The SNP distance matrix is standardized by R function scale () and then the function dist () (method=“euclidean”) is used to calculate the distance between variables. The results are visualized by using the function cmdscale and by Package ggplot2 ().

**Table 1.** Results of permanova variance analysis among clonalities based on SNP distance. (permutation = 10000)

| Group | SumsOfSqs | MeanSqs | F.Model | R<sup>2</sup> | Pr(>F) |
| --- | --- | --- | --- | --- | --- |
| Clone1/Clone2 | 13.8085505 | 13.8085505 | 270758.7184 | 0.99976 | 0.001*** |
| Clone1/Clone3 | 12.1801455 | 12.1801455 | 620304.5903 | 0.9999017 | 0.001*** |
| Clone1/Clone4 | 3.93263967 | 3.93263967 | 942.8360348 | 0.9505967 | 0.001*** |
| Clone1/Clone5 | 4.71278078 | 4.71278078 | 2737.589795 | 0.9820634 | 0.001*** |
| Clone1/Clone6 | 6.47426112 | 6.47426112 | 17532.50712 | 0.9970429 | 0.001*** |
| Clone2/Clone3 | 9.15172599 | 9.15172599 | 156083.8014 | 0.9997694 | 0.001*** |
| Clone2/Clone4 | 3.8765566 | 3.8765566 | 7205.820792 | 0.9966804 | 0.001*** |
| Clone2/Clone5 | 4.33333067 | 4.33333067 | 8269.488866 | 0.996986 | 0.001*** |
| Clone2/Clone6 | 5.50382535 | 5.50382535 | 13081.96483 | 0.9979403 | 0.001*** |
| Clone3/Clone4 | 3.84579693 | 3.84579693 | 1934040.787 | 0.9999897 | 0.001*** |
| Clone3/Clone5 | 4.34389168 | 4.34389168 | 673012.9105 | 0.9999688 | 0.001*** |
| Clone3/Clone6 | 5.40799762 | 5.40799762 | 2447588.684 | 0.9999906 | 0.001*** |
| Clone4/Clone5 | 2.24636341 | 2.24636341 | 1012.297445 | 0.9911877 | 0.002** |
| Clone4/Clone6 | 3.02713091 | 3.02713091 | 211478.7872 | 0.999948 | 0.002** |
| Clone5/Clone6 | 3.17394014 | 3.17394014 | 10754.74947 | 0.9988855 | 0.002** |
| Clone1/Clone other(B56) | 0.54775975 | 0.54775975 | 12.16291191 | 0.2127763 | 0.018* |
| Clone1/Clone other(A191) | 0.86621581 | 0.86621581 | 47.19376141 | 0.5118976 | 0.026* |
| Clone other(A191)/Clone2 | 0.90627404 | 0.90627404 | 131.2353356 | 0.8677558 | 0.044* |
| Clone2/Clone other(B56) | 0.64017946 | 0.64017946 | 12.43001057 | 0.3832873 | 0.044* |
| Clone3/Clone other(B56) | 0.93355985 | 0.93355985 | 4476.969255 | 0.9964389 | 0.053 |
| Clone other(A191)/Clone3 | 0.9409506 | 0.9409506 | 42274.54415 | 0.9996217 | 0.06 |
| Clone other(A191)/Clone6 | 0.87492897 | 0.87492897 | 2740.37143 | 0.9974521 | 0.115 |
| Clone6/Clone other(B56) | 0.77836812 | 0.77836812 | 64.50872318 | 0.9021098 | 0.115 |
| Clone5/Clone other(B56) | 0.26723539 | 0.26723539 | 2.371854408 | 0.3217446 | 0.135 |
| Clone other(A191)/Clone5 | 0.28517158 | 0.28517158 | 2.834401976 | 0.3617892 | 0.14 |
| Clone other(A191)/Clone4 | 0.81305707 | 0.81305707 | 3575.983694 | 0.9988827 | 0.166666667 |
| Clone4/Clone other(B56) | 0.74298762 | 0.74298762 | 212.3640798 | 0.9815126 | 0.166666667 |
| Clone other(A191)/Clone other(B56) | 0.5 | 0.5 | NaN | 1 | NA |

**Table 2.** Results of permanova analysis of variance among clusters (permutation = 10000)

| Group | SumsOfSqs | MeanSqs | F.Model | R2 | Pr(>F) |
| --- | --- | --- | --- | --- | --- |
| Cluster1/Cluster2 | 3.107777407 | 3.107777 | 47.80067 | 0.319095 | 0.001*** |
| Cluster1/Cluster3 | 0.488015237 | 0.488015 | 4.655904 | 0.051358 | 0.011* |
| Cluster2/Cluster3 | 0.933559854 | 0.93356 | 4476.969 | 0.996439 | 0.062 |

### Temporal distance identified relevant epidemiologic groups

We leveraged time, source, ages and gender information to identify epidemiologically meaningful groupings with the chi-square test (Fig 5). We found that the significant difference in community structure was only present in strains from separation time (p=2.149e^-11^), absent in sampling sources (p=0.2283), age of patients (p=0.0931) and gender of patients (p=0.0853). The significant difference in sampling time may be due to the changes in the main clone types in 2013. Furthermore, The clone 1 and clone 2 were all in high proportion of CRAB population across the whole collection time. We further analyzed the relationship between clone type and ICU rooms, the phenomenon of different clone types found in the same room was very common in hospital (Fig 6). Due to the hospital moved into a new place in 2018, we compared the difference of clone groups between the two hospitals, their main clone groups changed. The clone 6 and clone other (A191) were mainly found in 2018, while the clone 1 and clone 2 were still present in 2018.

**Fig 5.**
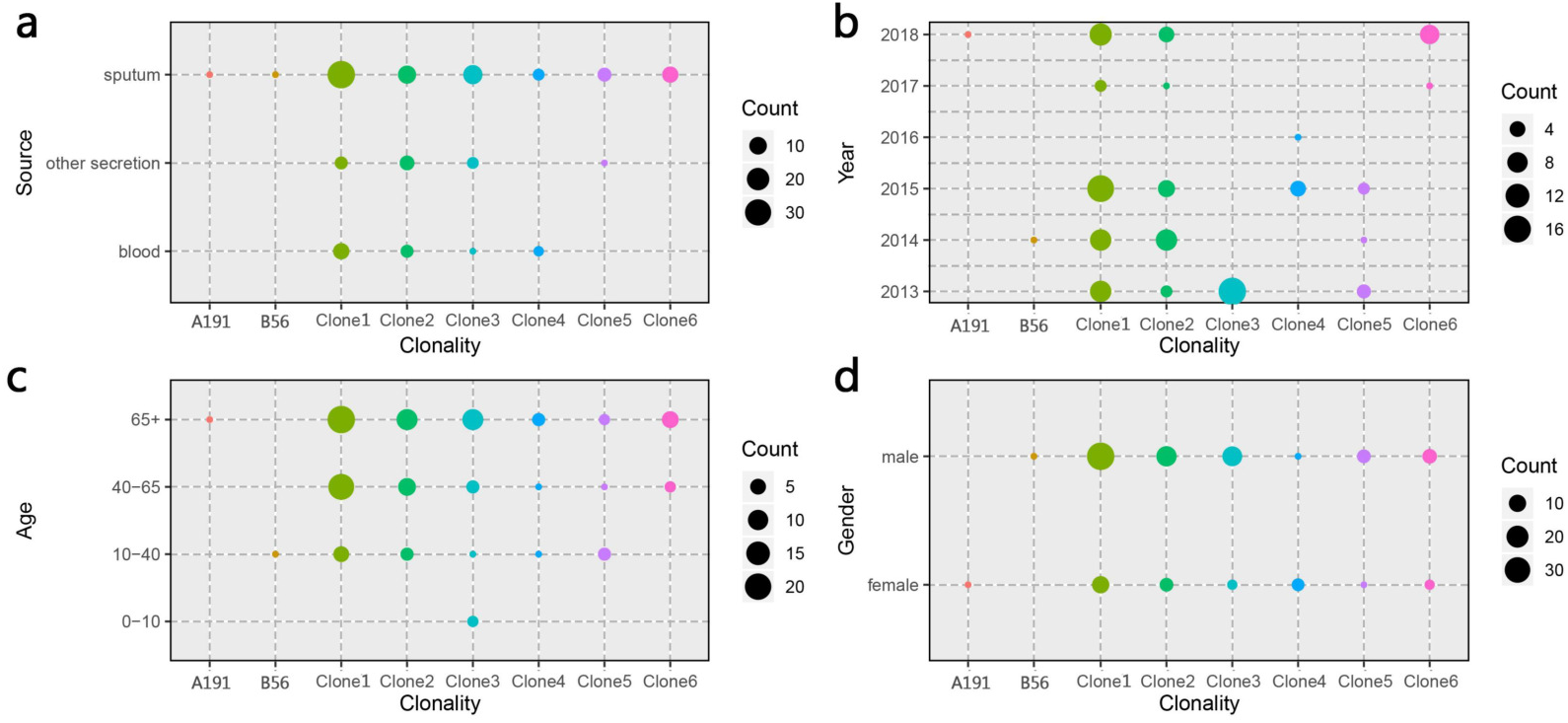
Distribution of CRAB clonalities in different spatial temporal, demographic and clinical characteristics. **a** Distribution of CRAB clonalities in different sampling source. **b** Distribution of CRAB clonalities in different sampling years. **c** Distribution of CRAB clonalities in different patient ages. **d** Distribution of CRABs clonalities in different patient genders.

**Fig 6.**
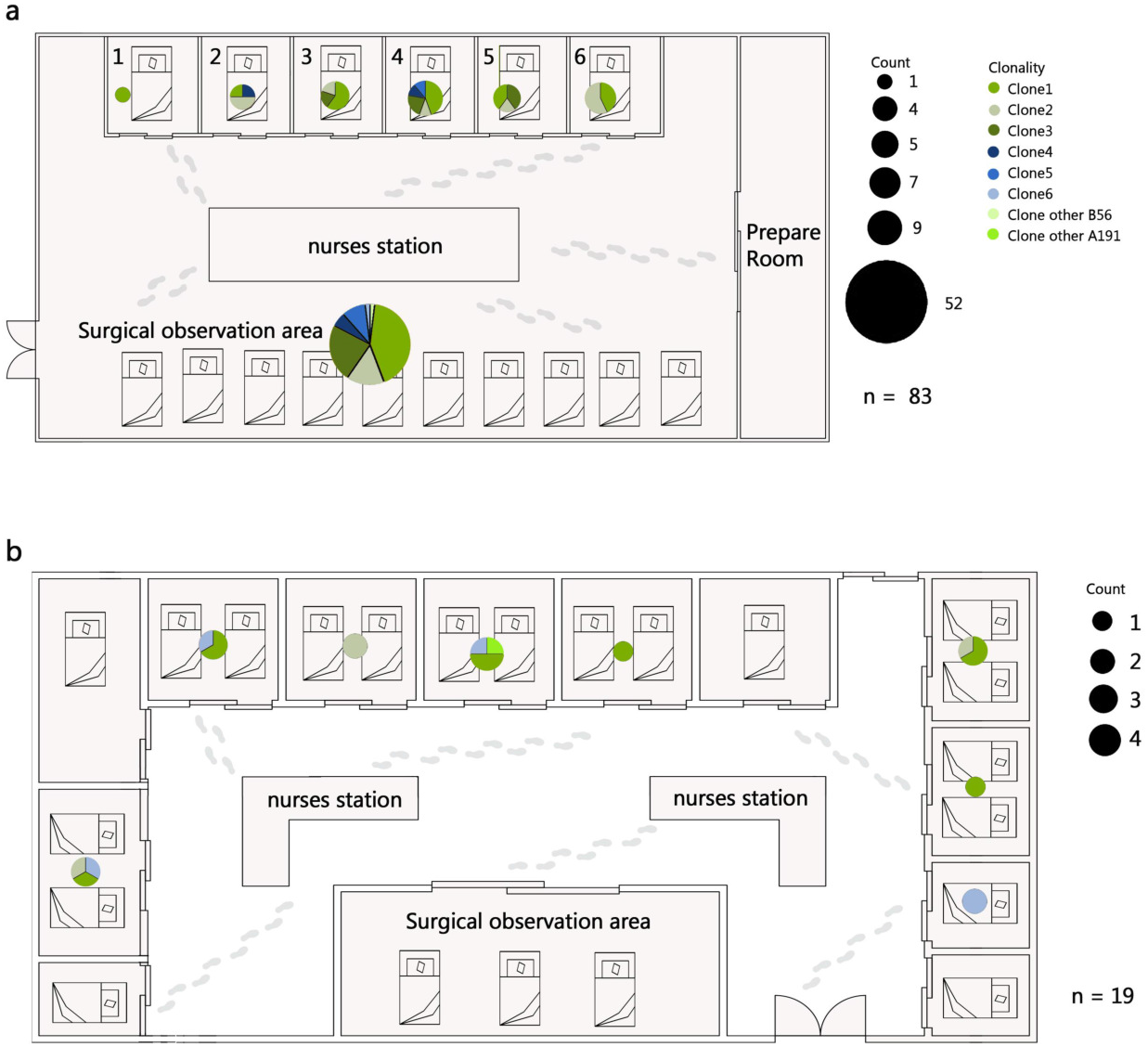
Distribution of CRAB clonalities in different ICU rooms. **a** Distribution of CRAB clonalities in different ICU rooms from 2013 to 2017. **b** Distribution of CRAB clonalities in different ICU rooms in 2018.

According to haplotype generated by DnaSP and the median-joining network generated by NETWORK based on SNP matrix calculated from core genome alignment of CRAB samples, these strains can be divided into 55 haplotypes. There were many haplotypes within close genetic relationship in the same clone type, and the haplotype genetic relationship between different clone groups was quite different, which is also accordant with our classification expectations. There were large genetic differences among different bayesian clusters. Over time, cluster 2 (only exists in 2013) gradually disappeared and cluster 1 subsequently became the dominant of the CRAB samples (Fig 7).

**Fig 7.**
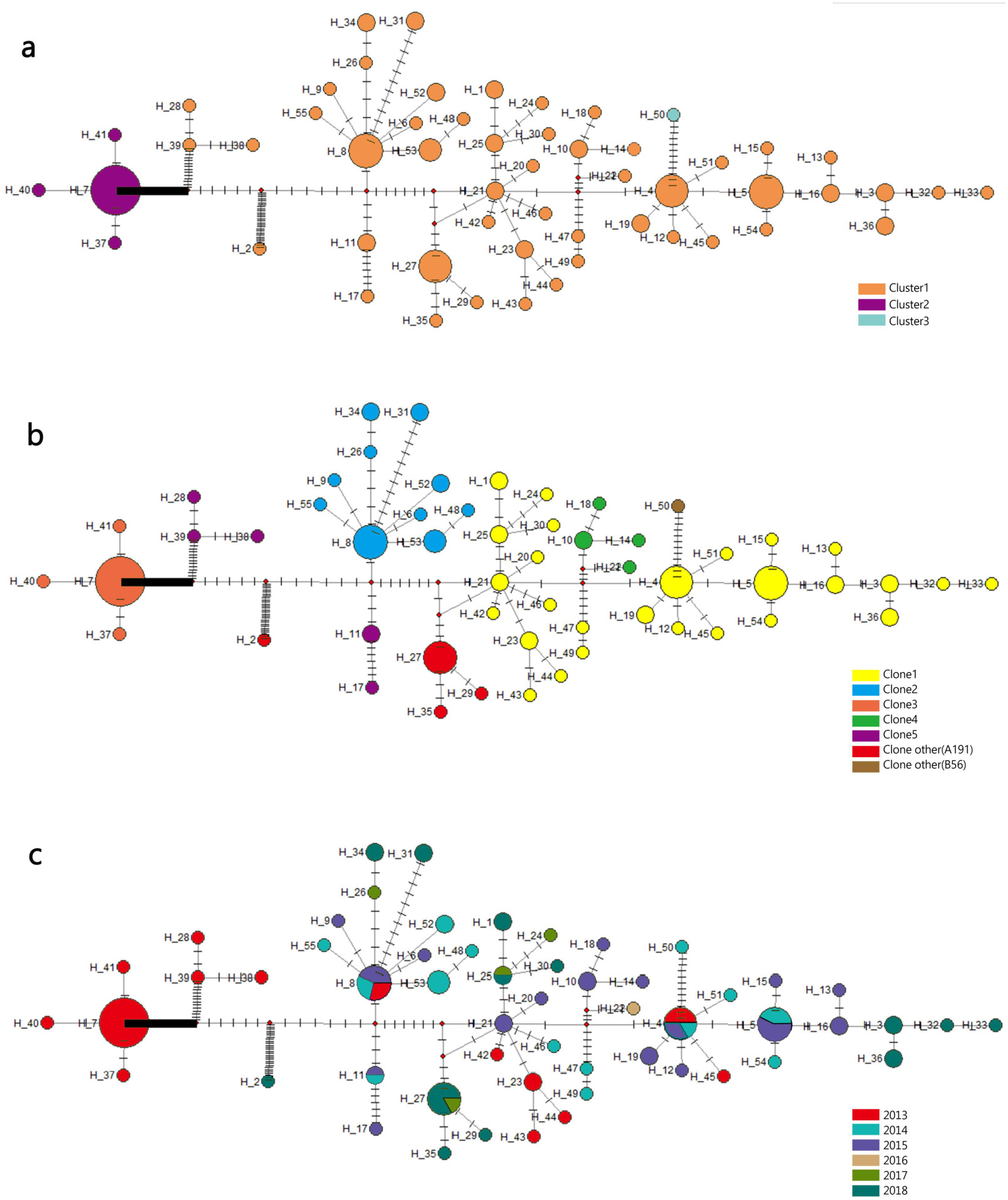
Evolutionary relationships and geographical distribution of 55 haplotypes of 105 isolates from Shunde hospital; circles size represent the number of isolates in a haplotype. **a** Circle colors represents different clusters based on bayesian classification. **b** Circle colors represents different clonalities based on SNP distance **c** Circle colors represents Sampling time. The density of those bars represents the genetic distance between the haplotypes

### Population expansion in ICU CRAB clusters

For more accurate results, we reconstructed the core genome alignment of 104 CRAB genomic after removing outliers B56 and M19606 for further population variation analysis (faced to cluster1 and cluster 2). According to the mismatch curve drawn by DnaSP, it can be found that the two main clusters were based on bayesian classification. The mismatch curve of total population (cluster1 and cluster 2) and cluster1 deviated from the expectation curve of assumed constant population size. That was also consistent with the results of Arlequin mismatch analysis, which was based on curve fitting and indicated that the confidence of fitting curve of total population (p = 1>>0.05) and cluster 1 (p=0.97>>0.05) (bootstrap = 1000) could not reject the population expansion hypothesis. Similar phenomenon also existed in the analysis of cluster 2 (p=0.27>0.05), but it was not obvious due to the problem of sample quantity (Fig 8). To sum up, it can be speculated that CRAB with different genetic backgrounds in ICU presented different degree of population expansion.

**Fig 8.**
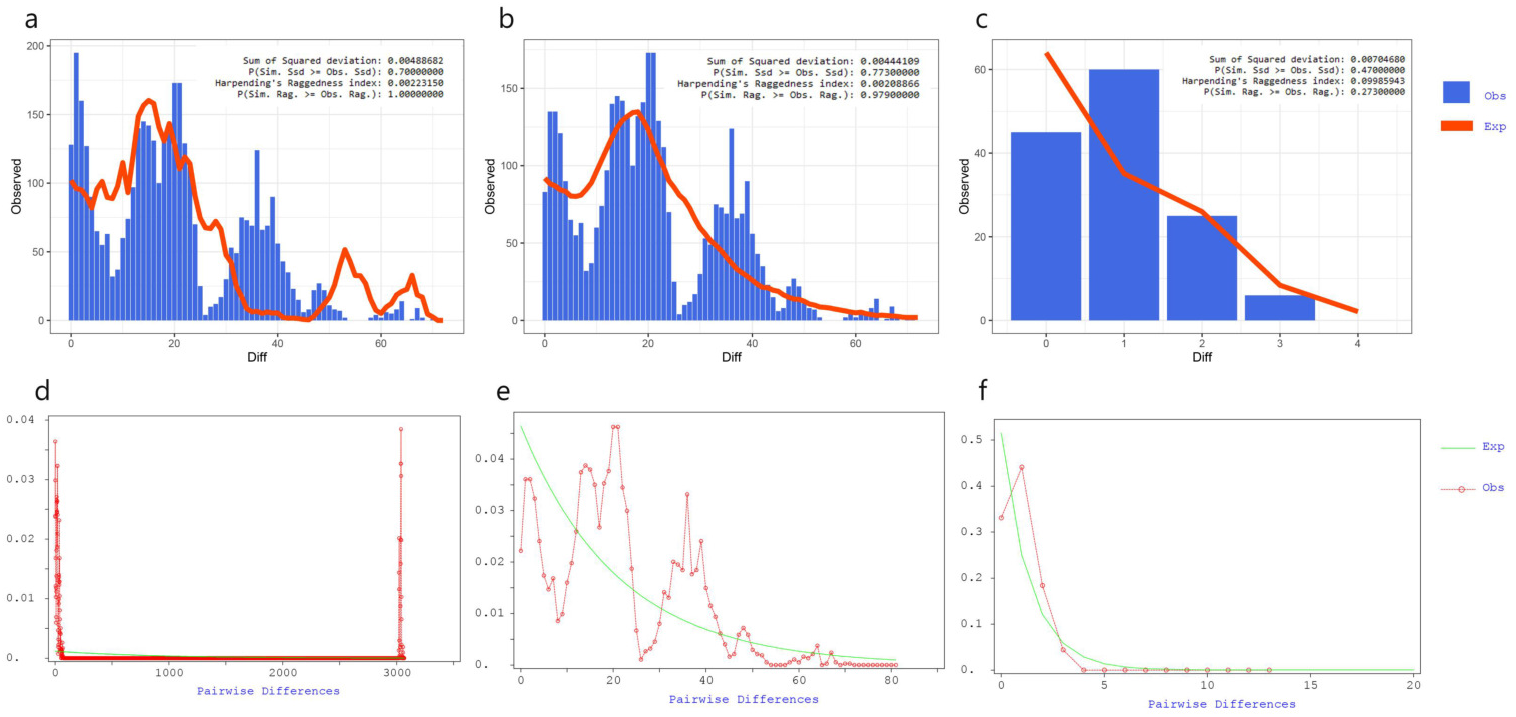
Mismatch distribution analyses of coding region alignment of 105 genomes from Shunde hospital. **a-c** Results of Mismatch distribution analyses by Alrequin. **d-f** Results of Mismatch distribution analyses by DnaSP.

### CRAB isolates shared high genotypic and phenotypic resistance, virulence factors and insertion sequences

Antibiotic susceptibility testing results revealed that all CRAB strains were resistant to at least three classes of antibiotics. All isolates showed 100% resistance against amoxicillin, ampicillin, cefoxitin, cefazolin, ceftazidime, gentamicin, chloramphenicol and florfenicol (Fig 9a). The rate of other resistances was as follows: 99.05% for ampicillin/sulbactam, cefotaxime, cefepime, ciprofloxacin, tetracycline, fosfomycin, streptomycin and piperacillin, 98.10% for aztreonam, levofloxacin and piperacillin/tazobactam, 92.38% for amikacin, 87.62% for ceforazone/ sulbactam, 79.05 for neomycin, 69.52% for trimethoprim/ sulfamethoxazole, 25.71% for tigecycline, and 12.38% for minocycline. None of them was resistant to polymyxins.

**Fig 9.**
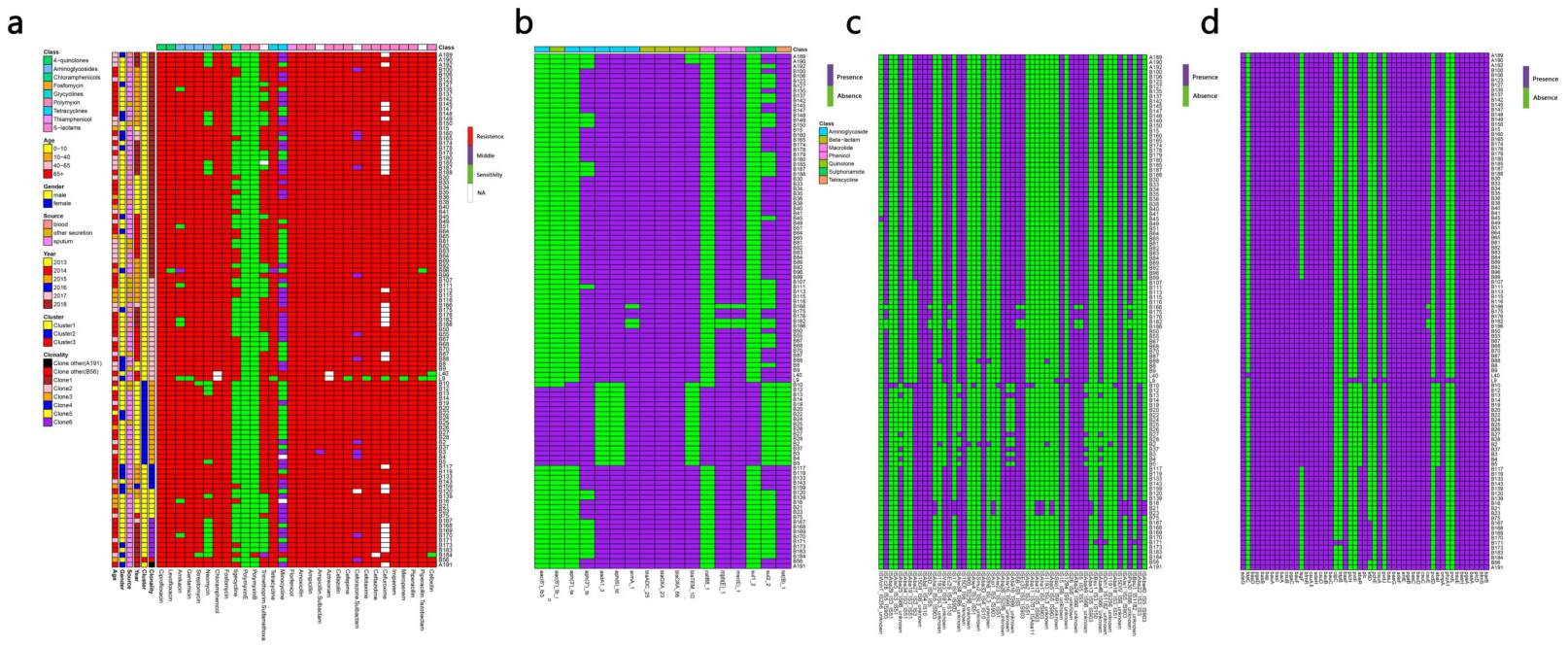
MIC test and ARGs, Virulence genes (VGs), insertion sequences (IS) identification of 105 samples from ICU rooms. **a** Results of MIC test; class of antibiotics and resistance strength are annotated as colored bars next to the main heatmap. **b** Results of ARGs identification based on Resfinder **c** Results of IS based on ISfinder **d** Results of VGs identification based on VFDB.

WGS analysis demonstrated that CRAB isolates harbored 17 unique ARGs. Carbapenemase genes *bla_OXA-23_, bla_OXA-66_* and cephalosporinase gene *blaAD_C-25_* were present in all strains. Genes encoding resistance to β-lactams (*bla_TEM-1D_*) aminoglycoside (*aac(6’)-Ib, aac(6’)Ib-cr, aph(3’)-Ia, aph(3’)-Ib, aph(6’)-Id, armA, aadA*), macrolide (*mph(E), msr(E)*), amphenicol (*catB8*), tetracycline (*tet(B)*) and sulphonamide ARGs (*sul1, sul2*) were also detected (Fig 9b).

Given the extensive burden of high-risk ARGs found in CRAB isolates, we analyzed the existence of VFs and ISs. We identified 55 differernt types of ISs and 51 unique VFs (Fig 8c and 8d). All the CRAB isolates had *bau (ABCDEF), bas (basABCD, basFGHIJ), csuABCD, pgaABCD, adeFGH, bfmRS, bap, entE, ompA*, and *plcD*. Interesting, there were three isolates carrying unique VFs. The *pvdLDJI, phzA, pchE, plcH* genes were only present in L9 strain, the *hptB* and *clpC* genes in B171 strain, and the *fdeC* in L40 strain. ISs screening in all genomes showed IS*Aba1/26/33*, IS*4*, IS*6*, *IS10*, IS*15*, IS*26*, IS*256* were found in all CRAB strains.

Besides, the hierarchical clustering of isolates based on ARGs, VFs and ISs presence indicated that cluster was the major predictor of resistance, virulence and IS patterns. Compared to cluster 1 and 3, ARGs like *aac(6’)-Ib, aac(6’)Ib-cr, aph(3’)-Ia, catB8, sul1* genes were nearly present in cluster 2strains, while the genes *aadA, aph(6’)-Id, bla_TEM-1D_, tet(B*) were almost absent. In addition, the strains in cluster 2 were more likely to lack the *AbaIR*, IS*Aba46*-IS*66*, IS*Aba49*-IS*66*, IS*Aba16*-IS*66*, IS*Aba25*-IS*66*, IS*Alw34*-IS*66*, IS*Alw16*-IS*66* and IS*Vsa3*-IS*91*.

## Discussion

CRAB is a perilous nosocomial pathogen causing substantial morbidity and mortality, which is a substantial health threat and economic burden(5, 25). Hospital setting is an important reservoir for CRAB transmission(41, 47). This study shed some light on the population dynamics of CRAB and demonstrated the endemicity and dissemination of clonal complex 2/92 (Pasteur scheme/oxford scheme) in the ICU.

Hospitalized patients with open wounds and long hospital stays are among those who are vulnerable to *A. baumannii* infections, especially those hospitalized in intensive care units (ICU)(26, 55). It was noted from this study that most patients were elder people, who had weak natural defenses and long hospital stays. Besides, CRAB isolates are responsible for a wide range of infections, including pulmonary infection, abdominal infection, bloodstream infection, septic shock and so on. Interestingly, almost all patients suffered from several diseases at the same time, the most common infection in this study was pulmonary infection, including pneumonia, respiratory failure. It was consistent with that CRAB infection were often associated with pneumonia and bloodstream infection(3, 38, 54).

Carbapenem resistance in *A. baumannii* is mainly based on the production of carbapenemases(23). In our study, all strains harbored the *bla_OXA-23_* and *bla_OXA66_* responsible for carbapenem resistance. This parallels previous investigations that *bla_OXA-23_* was the most widely reported gene and *bla_OXA66_* was the most common variant of OXA genes (also known as *bla_OXA51-like_*)(46). The increased antibiotic resistance in *A. baumannii* is largely owing to the actions of mobile genetic elements, activation of intrinsic resistance mechanisms such as the chromosomal β-lactamases, *blaADC* and the presence of efflux pumps(31). This study showed that a large diversity of ISs and AbaR-type genomic resistance islands (AbaRI) were present. It was noteworthy that all strains carried IA*Sba1/26/33*, IS*4*, IS*6*, IS*10*, IS*15*, IS*26* and IS*256*. Insertion of IS*Aba1* in the *bla_OXA-23_* promoter sequence has been reported to be associated with overexpression of *bla_OXA-23_, bla_OXA-51_* and causes carbapenem resistance phenotypes in *A. baumannii(32*). In addition, the presence of AdeFGH RND efflux pumps, which has been reported to play a major role in acquired resistance and may also be associated with carbapenem nonsusceptibility(10, 60). In accordance with others reports, these genetic structures could disseminate antibiotic resistance determinants between pathogens and thus compromise antibiotic treatment, including those prescribed antibiotics(1, 43). In this study, The CRAB isolates had a high ARG burden and were phenotypically resistant to multiple classes of antibiotics commonly used in clinical. Particularly problematic are the 28 CRAB isolates showing resistant to tigecycline, which limited treatment options for their infections.

It is largely believed that drug resistance and virulence factors have enabled *A. baumannii* to thrive in the unfavourable conditions, particularly in nosocomial environment(39). Interestingly, the virulome of CRAB is conserved in all CRAB strains. The presence of virulence factors, such as *bap*, *csuABCD*, *pgaABCD*, *bfmRS*, *entE*, *ompA* and *plcD* suggests the infectious property of these strains(8)’(35). The Csu pili (*csuA/BABCDE*, regulated by the BfmRS two component regulatory system), biofilm-associated proteins (*bap*), the production of polysaccharide poly-N-acetylglucosamine (*pgaABCD*) were proved to be implicated in the biofilm formation, maintenance and maturation(16, 29). Studies have found that biofilm formation is more strongly associated with MDR *A. baumannii* strains or clinical isolates. Besides, most of the acinetobactin biosynthetic genes (*basABCDFGHIJ*) or the genes involved in acinetobactin uptake (*bauABCDEF*), siderophore efflux system (*barAB*) were all present in CRAB isolates, which involved in the mechanism of iron uptake(22). The presence of the most important virulence genes in all CRAB isolates is in agreement with their clinical importance and confirmed the complexity of clinical infections caused by these strains.

There have been many reports of clonal outbreaks of *A. baumannii* mainly related to three international clone lineages (European clones I, II, and III)(61). For the MLST scheme, most outbreak strains belong to CC1 and CC2, corresponding to European clones I and II, respectively. So far, ST2 type as the most common ST in CC2 has been reported in many countries, including China. The results of this study showed that all CRAB isolates belonged to ST2, suggesting at least an outbreak situation in ICU rooms. Of note, *bla_OXA-23_* and *bla_OXA66_* positive ST2 isolates were responsible for many outbreak infections(24). Besides, ST2, ST25 and ST78 strains produced more biofilm than other STs, which was consistent with our strains carrying several virulence factors responsible for biofilm formation(45). This fact has potentially led to more successful colonization of the clinical environment over time.

Through core genome phylogenetic analysis, we found that our CRAB isolates are dominated by single lineage. Previous report of *A. baumannii* and *E. faecium* isolates from Pakistan and the USA hospital system showed they were similarly dominated by single lineages(11). Contrastingly, these strains were composed of three clusters and eight clones. A previous study reported that six of the eight Danish *A. baumannii* isolates were located on three distinct clusters based on core genome phylogenetic analysis(20). Two clusters have the strains assigned to ST195 and ST208, which both belonged to ST2. In our study, cluster 1 and cluster 2 were the main clusters, clone 1 and clone 2 were found in 5/6 timepoints during our six-year collections. This observation, particularly in the light of the time separation between isolation events, suggests existence of specific CRAB strains in the Shunde hospital and some predominant clones circulating in this North China region.

It was noted that changes in main clone groups were found in 2018 due to the hospital moved into another place. While the clone 1 and clone 2 were still found in 2018, which indicated that the clone outbreak may be associated with doctors and nurses or with the medical equipment. Especially, there were several clones coexisting in the surgical observation area of open environment, where patients stayed for one or two days after the operation. This increased the risk of cross infection among patients, as well as the main cause of nosocomial infection and bacteria outbreak. This phenomenon was improved evidently when increasing the number of inpatient wards and the closure of surgical observation area. These results aroused us to consider the effect of the hospital environment on nosocomial bacteria infections. Previous studies have reported on *A. baumannii* isolates from medical apparatus, water systems and handwashing sinks in the hospital environment, including ICUs and surgical wards(58).

According to the haplotype and the median-joining network analysis, we found large genetic differences among different clusters. Of note, cluster 2 was the dominant among all CRAB isolates at first (in 2013) and then substituted by cluster 1 between 2014 and 2018. These two clusters also presented different degree of population expansion. There was no study focused on the population variation analysis of the same ST CRAB strains. Isolates in cluster 1 and cluster 2 found to possess obvious different resistance, virulence and IS patterns further confirmed the genetic differences between the two clusters. Compared to cluster 1, cluster 2 contained relatively little IS and resistance gene types. This might led to the subsequent prevalent of cluster 1 in Shunde hospital. The origin of these clusters is unknown, but it seems likely that they dwell in the hospital environment and cause infections over an extended period of time. These isolates tended to be MDR and hence could persist for a long time if the reservoir is not destroyed, serving as a potential source for nosocomial infection and recurrent outbreaks. In conclusion, based on the molecular epidemiology and genomics data, we confirm that polyclonal nature of the CRAB outbreak in hospital mainly caused by ST2 clone carrying *bla_OXA-23_* and *bla_OXA-66_*.

## Conclusion

We found the wide dissemination and clone outbreak of CRAB ST2 in ICU rooms of hospital. All strains had high genotypic and phenotypic resistance burden and were dominated by a single lineage of three clusters and eight different clones. The main cluster changed over years and CRAB strains with different genetic backgrounds presented different degree of population expansion.

## Author contributions

W-B L and Z-L Z conceived this study and designed the experiments.

X-F Z and F-P L drafted the manuscript. W-B L offered all the CRAB strains. X-F Z, F-P L, Z-W H, F A performed the bioinformatic analysis. H-Y J performed antimicrobial susceptibility tests for all CRAB strains. X-H L made the data visualized. All authors read and approved the final manuscript.

